# Promotion of seed germination in *Echinochloa crus-galli* with atmospheric air plasma

**DOI:** 10.1101/2025.11.14.688452

**Authors:** Yuto Nakamura, Yu Ohnishi, Kazuma Oshimo, Rei Hiramatsu, Masako Shindo, Tohru Tominaga, Satoshi Terabayashi, Daisuke Takagi, Mitsuhiro Matsuo

**Author notes:** Author for correspondence: **Mitsuhiro Matsuo;**.

## Abstract

Plasma technology has attracted increasing attention for potential agricultural applications, and it has been investigated for decades particularly for its ability to promote seed germination. However, most studies have focused on crop species, and little is known about its potential use for weed management. In this study, we examined whether air plasma influences the germination of *Echinochloa crus-galli*, a major paddy field weed. When seeds were cultivated without stratification, direct irradiation with air plasma did not enhance germination. In contrast, once dormancy was partially released by cold stratification, plasma treatment significantly promoted germination. Interestingly, the effect disappeared after prolonged chilling, when dormancy was completely released. These results suggest that plasma promotes germination by acting on the dormancy-breaking process. In addition, plasma-activated water (PAW) also stimulated germination, although its effect was weaker than that of gaseous plasma. Overall, this study demonstrates that air plasma can promote the germination of grass weed seeds, providing a potential strategy for weed control by inducing germination prior to crop seeding.

## Introduction

Weeds are considered major pests that significantly reduce crop yields (Chauhan, 2020; Oerke, 2006). To maintain high levels of crop productivity, effective weed management strategies are essential. A key trait that leads to the persistence of weeds and increases the burden on farmers is their year-round emergence. This trait is primarily attributed to seed dormancy, seed longevity and responsiveness to environmental changes. Dormancy serves as an adaptive mechanism that prevents germination during unfavorable seasons or under suboptimal environmental conditions (Finch Savage and Leubner Metzger, 2006). During this period, seeds remain metabolically inactive yet viable. In addition, many weed seeds exhibit long-term viability, persisting in the soil seedbank for several years or even decades (Conn et al., 2006). These seeds may germinate opportunistically when environmental conditions become favorable, complicating weed control and increasing labor and management costs.

To mitigate the difficulty of weed management, the stale seedbed technique has been developed as a cultural weed management strategy (Gallandt, 2006; Travlos et al., 2020). This method promotes weed seed germination before crop sowing through soil disturbance practices such as shallow tillage. It exploits the tendency of many weed seeds to germinate in response to light exposure, temperature fluctuations, and moisture availability. Once germinated, these seedlings can be eliminated by mechanical, thermal, or chemical means prior to planting the main crop, thereby reducing weed pressure during the critical early stages of crop development. Furthermore, in the case of *Striga* spp. (witchweed), which are parasitic weeds of major concern in sub-Saharan

Africa, synthetic strigolactones, which are compounds that stimulate *Striga* germination, can be applied in the absence of a host plant (Zwanenburg et al., 2016). As a result, *Striga* seeds germinate and subsequently die due to the lack of a suitable host. This “suicidal germination” approach is conceptually similar to the stale seedbed technique and highlights the broader potential of exploiting seed biology in weed management. These approaches not only reduce the frequency of weeding during crop cultivation but also help deplete the weed seedbank in the soil. Continued development and refinement of such techniques are essential for sustainable agricultural systems.

In recent decades, the application of plasma technology to plant science and agriculture has been intensively studied (Bourke et al., 2018). One important application is seed processing technology, in which plasma treatment is used to promote seed germination (Grainge et al., 2022; Waskow et al., 2021). Plasma treatment has been suggested to induce alterations in the physical and biochemical properties of seeds, thereby influencing germination (Waskow et al., 2021). In particular, air plasma abundantly generate reactive oxygen species (ROS) and reactive nitrogen species (RNS), which are considered as key agents because they have been reported as signaling molecules regulating seed germination (Bykova and Igamberdiev, 2024; Puglia, 2024). The positive effects of air plasma on seed germination have been reported across a wide range of species, including not only angiosperms but also gymnosperms (Priatama et al., 2022; Waskow et al., 2021). However, most studies have focused on crop species, whereas knowledge related to weed management remains limited. Bukhori et al. (2024) reported that high-voltage plasma treatment reduced the germination rate of mustard seeds and suggested that plasma technology could potentially be applied to control weed seed germination. However, their study did not directly involve weed species and did not examine the potential application of the promotive effects of plasma on seed germination in the context of weed control.

Theoretically, promoting germination with plasma treatment, followed by the elimination of seedlings before crop planting, could be an effective strategy for weed control and soil seedbank management. In this study, we investigated the effect of plasma on the germination of *Echinochloa crus-galli*, one of the most problematic weeds in both paddy and upland fields (Tian et al., 2020; Zhang et al., 2021), and showed that plasma could induce the germination in a manner dependent on dormancy status. Our findings suggest that plasma could serve as a novel tool to stimulate weed seed germination for pre-sowing weeding and soil seedbank regulation.

## Material and methods

### Plant material and cultivation

Seeds of *Echinochloa crus-galli* were collected from a paddy field in Higae-cho, Keihoku, Ukyo-ku, Kyoto City, Japan (35.1819°N, 135.6665°E). The collected seeds were propagated in a controlled-environment growth chamber to increase the seed stock. Seeds obtained from the cultivations were used for the germination experiments. Plants were grown under controlled conditions of 25 °C with a 14-hour light and 10-hour dark photoperiod.

### Plasma irradiation and construction of plasma-activated water

Atmospheric pressure air plasma was generated using a dielectric barrier discharge (DBD) system with pen-type plasma device. The system was constructed of power supply (LHV-13AC, LOGY ELECTRIC CO.,LTD., Tokyo, Japan), an air-pump (e-air 2000SB, Gex, Osaka, Japan) and a handmade pen-type electrode. The electrode was constructed by placing a stainless wire mesh inside a ceramic tube (4 mm inner diameter, 6 mm outer diameter) as the high-voltage electrode, and attaching a copper sheet on the outside of the ceramic tube as the grounded electrode. The air plasma was applied to the seeds of *Echinochloa crus-galli* for 15 minutes. Plasma-activated water (PAW) was prepared by introducing the plasma into 1 L of deionized water for 3 hours at 22 °C, with a flow rate of 0.5 L/min. Deionized water aerated without plasma for 3 hours were used as control. PAW and control water was applied to the double ring filter paper (99-191-090, Cytiva, USA), where the seeds were sown.

### Germination test

30 seeds were sown on filter paper. Stratification of the seeds were performed at 4 C°C under darkness for the specified periods. And then the they were transferred to light condition at 22 °C. Germinated seeds were counted and the average germination ratio and standard deviation were calculated from three independent experiments.

### Measurements of nitrate, nitrite and hydrogen peroxide

NOCC and NO_2_C were measured using the spectroscopic method described by according to (Hachiya and Okamoto, 2017). H_2_O_2_ was measured using Amplite®Fluorimetric Hydrogen Peroxide Assay Kit (Red fluorescence, AAT Bioquest, Pleasanton, CA. USA) according to the manufacture’s instruction.

## Results

### Pen-type DBD plasma irradiation system and the Plasma-activated water

In this study, air plasma was generated using a handmade pen-type dielectric barrier discharge (DBD) plasma system and used for seed treatment experiments. The system delivered air (0.7 L minC¹) into the nozzle electrode via an air pump, where it was converted into air plasma by applying a high voltage (2.3 W, Vpp 9 kV, 13.7 kHz) generated from an AC power supply (Figure 1A, B). The generated plasma, which emitted purple light (Figure 1C), was either directly irradiated onto *Echinochloa crus-galli* seeds or bubbled into water to prepare plasma-activated water (PAW) (Figure 1B).

**Figure 1.**
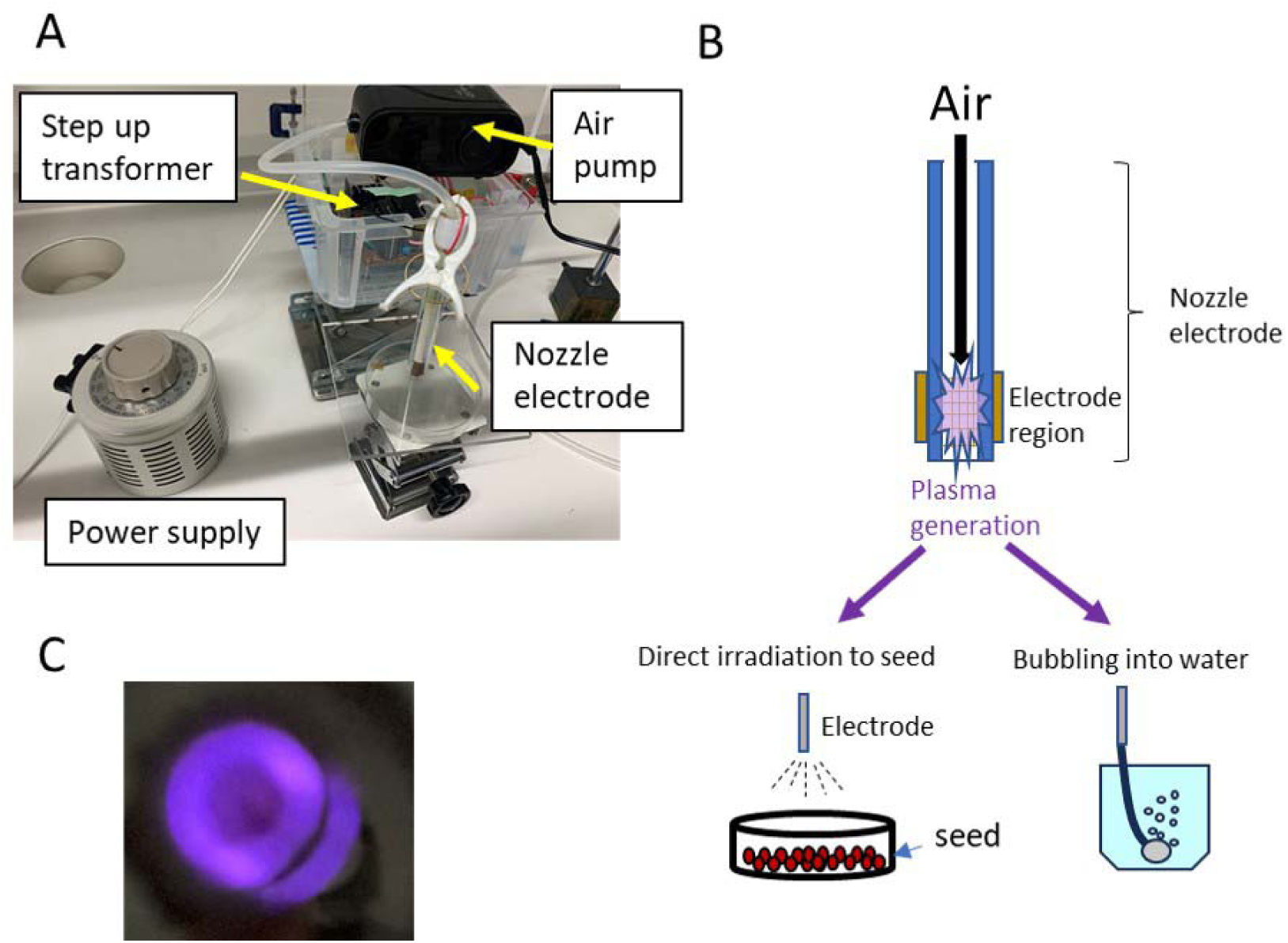
Generation of atmospheric air plasma and plasma-activated water (PAW). A) Overview of the handmade atmospheric air plasma generation device. The device mainly consists of four parts: a power supply, an air pump, and a nozzle electrode. B) Schematic illustration of the air plasma treatment. Air delivered by the pump to the nozzle electrode is ionized under high voltage to generate plasma. The ejected plasma was either directly irradiated onto *Echinochloa crus-galli* seeds or bubbled into deionized water to produce plasma-activated water (PAW). C) Air plasma emitting purple light inside the nozzle electrode.

### Plasma irradiation promotes the germination of *Echinochloa crus-galli* dependent on dormancy break

*Echinochloa crus-galli* seeds have deep dormancy, which can be broken by cold treatment (Honek et al., 1999). To explore the effect of air plasma on the germination of *Echinochloa crus-galli* seeds, seeds were treated with air plasma and incubated either with or without cold treatment (Figure 2A–D). The air plasma treatment significantly promoted germination when seeds were stratified with one month of cold treatment (Figure 2C). However, no promotive effect of air plasma was observed in seeds incubated without cold treatment or with only a short, one-week cold treatment, which was not sufficient to break dormancy and resulted in a low germination rate (Figure 2A, B). Furthermore, most seeds were germinated after two months of cold stratification and there was less difference between the plasma treated seeds and the control seeds (Figure 2D). These results suggest that air-plasma should promote the germination of *Echinochloa crus-galli* but the positive effect should occur during the periods when the dormancy is released.

**Figure 2.**
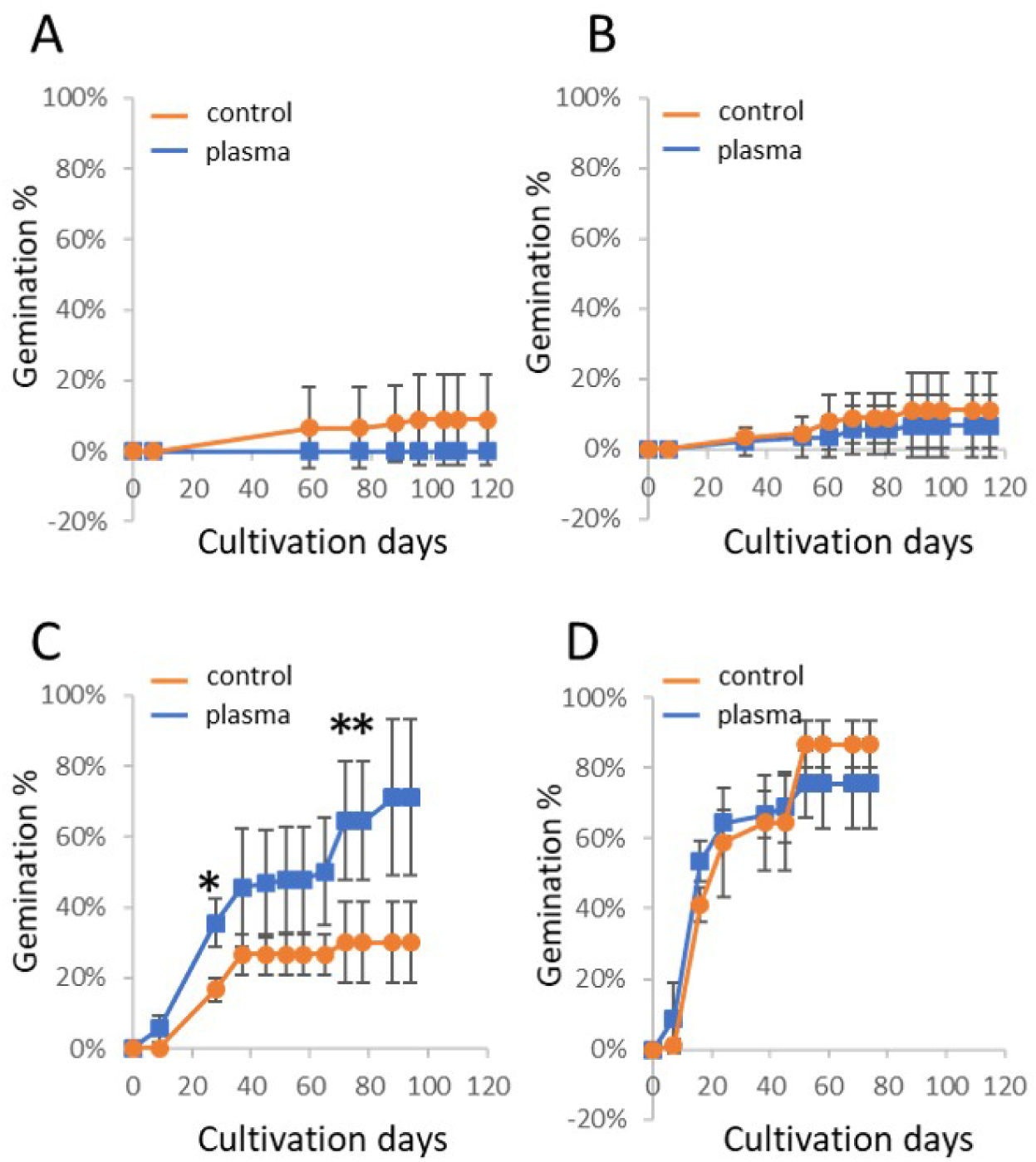
Effect of plasma irradiation on the germination of *Echinochloa crus-galli Echinochloa crus-galli* seeds were directly irradiated with atmospheric air plasma, and then sown on wet filter paper. Plasma-irradiated and control seeds were transferred from stratification conditions (4 °C, darkness) to germination conditions (22 °C, continuous light), and then cultivated for 2–4 months. The horizontal axis indicates the cultivation period under germination conditions, and the vertical axis represents the percentage of germinated seeds. A) No stratification. B–D) Seeds stratified for 1 week (B), 1 month (C), and 2 months (D), respectively. Data are presented as means ± standard deviations. Asterisks indicate significant differences according to the *t*-test (*p* < 0.05).

### Plasma-activated water promote the germination of *Echinochloa crus-galli*

When considering the application of the germination-promoting effect of plasma in agriculture, it would be practical to use air plasma in the form of plasma-activated water (PAW), produced by infusing air plasma into water. We examined whether PAW has a similar germination-promoting effect (Figure 3). When the seeds were placed on filter paper soaked in PAW, no promotion of the germination was observed in dormant seeds without cold treatment (Figure 3A). After 2 weeks cold treatment, in a part of the PAW treated seeds, germination was promoted (Figure 3B), although most of the seeds remained to be ungerminated. In contrast to direct plasma irradiation of Figure 2C, there was less promotion in 1 month cold treated seeds (Figure 3C). For 2 months cold treated seed, whose dormancy should have been broken, no promotion effect by PAW was detected as well as Figure 2D. These results suggest that the liquid form of air plasma should have germination promotion effect of *Echinochloa crus-galli* seeds during seed dormancy breaking periods. However the effect should be less compared to the direct plasma irradiation.

**Figure 3.**
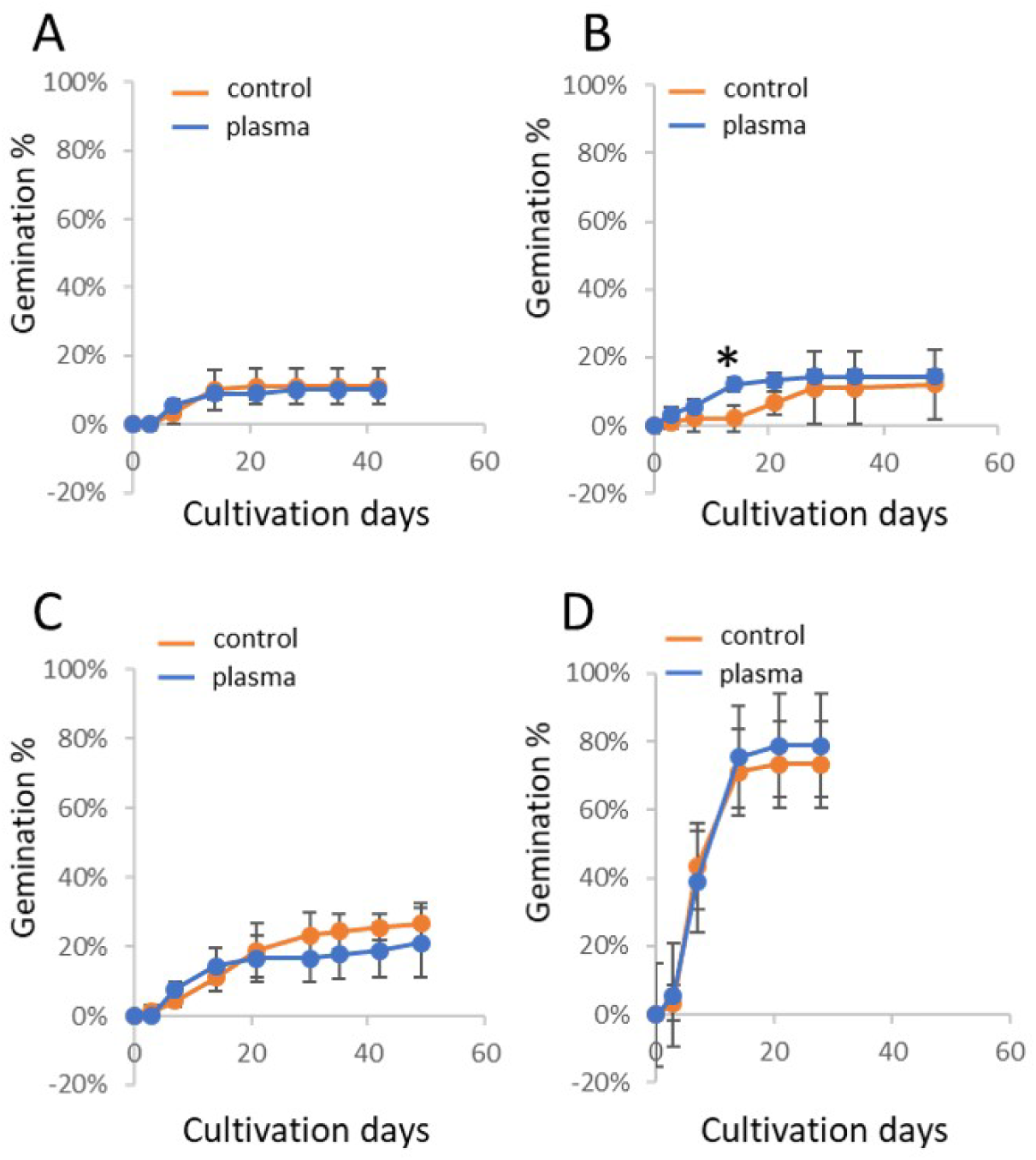
Effect of plasma-activated water (PAW) on the germination of *Echinochloa crus-galli Echinochloa crus-galli* seeds were treated with PAW or air-bubbled water (control) and then sown on wet filter paper. PAW-treated and control seeds were transferred from stratification conditions (4 °C, darkness) to germination conditions (22 °C, continuous light), and then cultivated for more than one month. The horizontal axis indicates the cultivation period under germination conditions, and the vertical axis represents the percentage of germinated seeds. A) No stratification. B–D) Seeds stratified for 2 weeks (B), 1 month (C), and 2 months (D), respectively. Data are presented as means ± standard deviations. An asterisk indicates a significant difference according to the *t*-test (*p* < 0.05).

### Plasma-activated water contain nitrate and hydrogen peroxide

It was reported that PAW of air could contain nitrate (NO_3_^-^), nitrite (NO_2_^-^) and hydrogen peroxide (H_2_O_2_) (Sivachandiran and Khacef, 2017). These molecules have been shown to have the effect of germination promotion (Bethke et al., 2006; Duermeyer et al., 2018; Wojtyla et al., 2016). To unveil how much these molecules are included in the PAW generated by pen-type plasma device in this study, we measured the amount of NO_3_^-^, NO_2_^-^ and H_2_O_2_ in the PAW. The analysis showed that the PAW contained 1.9 ×10^2^ μM NO_3_^-^ and 9.4 ×10^-1^ μM H_2_O_2_, but no NO ^-^ was detected (Table 1).

**Table 1.** Concentrations of NO_3_^-^, NO_2_^-^ and H_2_O_2_ in Plasma-activated water (PAW) PAW was prepared as described in Material and Methods section, and air-bubbled deionized water was used as control. nd, below detection limit. Data are presented as means ± standard deviations. n ≥ 3. Asterisk indicates significant differences compared with control according to *t*-test (p < 0.05).

| | $\text{NO}_3^-$ | $\text{NO}_2^-$ | $\text{H}_2\text{O}_2$ |
| --- | --- | --- | --- |
| | $\mu\text{M}$ | | |
| Control | nd | nd | $3.18 \pm 2.47 \times 10^{-1}$ |
| PAW | $1.90 \pm 0.80 \times 10^2$ | nd | $9.36 \pm 3.15 \times 10^{-1}$* |

## Discussion

It has been reported that atmospheric air plasma has positive effect on plant germination and the characteristics is expected to apply to agriculture. However there is few knowledge for utilization of the positive effect into weeding. In this study, we examined whether air plasma has germination promotion effect on *Echinochloa crus-galli*, one of the major weed in paddy field world wide, and showed that the plasma and plasma-activated water has the germination promotion effect in this weed.

Similar to many other weed species, *Echinochloa crus-galli* seeds exhibit long-term dormancy, and their germination largely depends on the degree of dormancy. To examine the effect of air plasma on germination and dormancy, we treated the seeds with the plasma and evaluated germination after different periods of cold stratification. Our results showed that the promotive effect on germination was evident only during the dormancy-breaking period in *Echinochloa crus-galli* (Figure 2C). In addition, no promotion was observed in seeds without cold stratification (Figure. 2A), and the effect disappeared once dormancy was fully released (Figure 2D). These results indicate that the plasma alone does not fully break dormancy but rather accelerates the dormancy-breaking process in *Echinochloa crus-galli*. Seed dormancy maintenance and release are regulated by highly complex molecular systems (Bykova and Igamberdiev, 2024; Puglia, 2024; Sajeev et al., 2024). Air plasma may act on regulatory factors to attenuate seed dormancy in *Echinochloa crus-galli*.

In this study, we showed that plasma-activated water (PAW) also promoted seed germination in *Echinochloa crus-galli*, but its effect was markedly weaker than that of direct plasma irradiation on seeds. This may be because the amount of germination-activating molecules in PAW is insufficient. Atmospheric air plasma is known to contain various reactive nitrogen and oxygen species (RNS and ROS) in abundance (Cimerman and Hensel, 2023). Among the RNS, nitogen oxide (NO) has been reported to promote seed germination as signal molecule (Arc et al., 2013). NO appears to be major factor responsible for the germination-promoting effect of air plasma. However NO is unstable and is rapidly oxdized into NO_2_^-^ and NO_3_ ^-^ in water. NO_3_^-^ also promotes germination and is likely one of key molecules regulating germination in PAW. Huang et al (2025) reported that the molecular mechanism by which NO_3_^-^ induces germination is distinct from that of NO (Huang et al., 2025). In addition, Bethke et al. (2006) demonstrated that a much higher concentration of NO_3_ ^-^ is required to break seed dormancy compared with NO in *Arabidopsis thaliana*. They showed that 100 ppm (≈4 μM) NO could induce about 40% seed germination, whereas approximately 1.0 mM NO_3_^-^ was needed to achieve similar germination rate (Bethke et al. 2006). In this study, 0.2 mM NO_3_^-^ was detected in PAW. This concentration is far lower than the NO_3_^-^ concentration, which induce germination, previously reported in *Arabidopsis thaliana* (Bethke et al. 2006) and *Echinochloa* colona (Picapietra and Acciaresi, 2022). The weak germination-promoting effect of PAW may be attributed to its low NO_3_^-^ concentration. In addition to NO_3_ ^-^, H O, which is one of the stable ROS in water, was also detected in PAW (Table 1), and it has been reported to promote seed germination (Ogawa and Iwabuchi, 2001). However, the concentration (0.9 μM, Table 1) is about two order of maginitude lower than that reported in other studies (Barba-Espín et al., 2012; Ogawa and Iwabuchi, 2001). Therefore the limitted effect of PAW may be due to the low concentration of these molecules. Increasing the concentration of NO_3_^-^ and H_2_O_2_ concentration by optimizing the PAW generation system may enhance its germination-promoting effect, although this remains to be investigated in future work.

The stale seedbed technique, which promotes weed germination and subsequent weeding, is widely used in agricultural practice, and its improvement could contribute to future sustainable agriculture. Our results showed that the germination of a grass weed could be promoted by plasma and PAW, suggesting that plasma technology has the potential to enhance and extend chemical-based stale seedbed techniques for weed management. Notably, the germination-promoting effect of plasma has been reported across a wide range of plant species (Priatama et al., 2022; Waskow et al., 2021). Its application to weeding does not seem to be limited to *Echinochloa crus-galli*, but could potentially cover diverse species. Furthermore, air plasma and PAW can be generated from air, water and household electricity using a simple device composed of a power supply, air pump and pen-type electrode. The simplicity of this device could enable on-site production of plasma and PAW, rather than relying on an industrial setting. In addition, plasma generates multiple molecular species, including NO, NOCC, and HCOC, which promote germination in a single step. The composition of air plasma can be adjusted depending on the plasma device and the generation conditions (Cimerman and Hensel, 2023; Klenivskyi et al., 2024). This flexibility allows the plasma composition to be tailored according to seed species, physiological state, and environmental conditions for agricultural applications. These characteristics may offer new alternatives in agricultural practice. However, the effects of plasma on germination and plant growth also depend on seed age and physiological state (Attri et al., 2021). Under natural field conditions, seeds in the soil exhibit diverse physiological states. Therefore, numerous factors need to be considered when applying plasma and PAW treatments to weed seed germination. Further investigations integrating plasma technology with plant science and agronomy are required to assess the practical feasibility and potential applications of plasma technology for weed control.

## Acknowledgement

This study was supported by JSPS KAKENHI Grant Numbers JP23K03371. We gratefully thank Prof. Junichi Obokata and Prof. Junshi Yazaki for their critical and productive discussions.

## References

Arc, E., Galland, M., Godin, B., Cueff, G., Rajjou, L., 2013. Nitric oxide implication in the control of seed dormancy and germination. Front. Plant Sci. 4. 10.3389/fpls.2013.00346

Attri, P., Ishikawa, K., Okumura, T., Koga, K., Shiratani, M., Mildaziene, V., 2021. Impact of seed color and storage time on the radish seed germination and sprout growth in plasma agriculture. Sci. Rep. 11, 2539. 10.1038/s41598-021-81175-x

Barba-Espín, G., Hernández, J.A., Diaz-Vivancos, P., 2012. Role of H_2_ O_2_ in pea seed germination. Plant Signal. Behav. 7, 193–195. 10.4161/psb.18881

Bethke, P.C., Libourel, I.G.L., Jones, R.L., 2006. Nitric oxide reduces seed dormancy in Arabidopsis. J. Exp. Bot. 57, 517–526. 10.1093/jxb/erj060

Bourke, P., Ziuzina, D., Boehm, D., Cullen, P.J., Keener, K., 2018. The Potential of Cold Plasma for Safe and Sustainable Food Production. Trends Biotechnol. 36, 615–626. 10.1016/j.tibtech.2017.11.001

Bukhori, A., Guntoro, D., Sudradjat Sugiarto, A.T., 2024. Utilization of Plasma Technology to Control Weed Seed Germination. J. Trop. Crop Sci. 11, 200–205. 10.29244/jtcs.11.02.200-205

Bykova, N.V., Igamberdiev, A.U., 2024. Redox Control of Seed Germination is Mediated by the Crosstalk of Nitric Oxide and Reactive Oxygen Species. Antioxid. Redox Signal. ars.2024.0699. 10.1089/ars.2024.0699

Chauhan, B.S., 2020. Grand Challenges in Weed Management. Front. Agron. 1, 3. 10.3389/fagro.2019.00003

Cimerman, R., Hensel, K., 2023. Multi-hollow Surface Dielectric Barrier Discharge: Production of Gaseous Species Under Various Air Flow Rates and Relative Humidities. Plasma Chem. Plasma Process. 43, 1411–1433. 10.1007/s11090-023-10381-4

Conn, J.S., Beattie, K.L., Blanchard, A., 2006. Seed viability and dormancy of 17 weed species after 19.7 years of burial in Alaska. Weed Sci. 54, 464–470. 10.1614/WS-05-161R.1

Duermeyer, L., Khodapanahi, E., Yan, D., Krapp, A., Rothstein, S.J., Nambara, E., 2018. Regulation of seed dormancy and germination by nitrate. Seed Sci. Res. 28, 150–157. 10.1017/S096025851800020X

FinchCSavage, W.E., LeubnerCMetzger, G., 2006. Seed dormancy and the control of germination. New Phytol. 171, 501–523. 10.1111/j.1469-8137.2006.01787.x

Gallandt, E.R., 2006. How can we target the weed seedbank? Weed Sci. 54, 588–596. 10.1614/WS-05-063R.1

Grainge, G., Nakabayashi, K., Steinbrecher, T., Kennedy, S., Ren, J., Iza, F., Leubner-Metzger, G., 2022. Molecular mechanisms of seed dormancy release by gas plasma-activated water technology. J. Exp. Bot. 73, 4065–4078. 10.1093/jxb/erac150

Hachiya, T., Okamoto, Y., 2017. Simple Spectroscopic Determination of Nitrate, Nitrite, and Ammonium in Arabidopsis thaliana. BIO-Protoc. 7. 10.21769/BioProtoc.2280

Honek, Martinkova, Jarosik, 1999. Annual cycles of germinability and differences between primary and secondary dormancy in buried seeds of *Echinochloa crus*lJ*galli*. Weed Res. 39, 69–79. 10.1046/j.1365-3180.1999.00122.x

Huang, Z., Han, X., He, K., Ye, J., Yu, C., Xu, T., Zhang, J., Du, J., Fu, Q., Hu, Y., 2025. Nitrate attenuates abscisic acid signaling via NIN-LIKE PROTEIN8 in Arabidopsis seed germination. Plant Cell 37, koaf046. 10.1093/plcell/koaf046

Klenivskyi, M., Khun, J., Thonová, L., Vaňková, E., Scholtz, V., 2024. Portable and affordable cold air plasma source with optimized bactericidal effect. Sci. Rep. 14, 15930. 10.1038/s41598-024-66017-w

Oerke, E.-C., 2006. Crop losses to pests. J. Agric. Sci. 144, 31–43. 10.1017/S0021859605005708

Ogawa, K., Iwabuchi, M., 2001. A Mechanism for Promoting the Germination of Zinnia elegans Seeds by Hydrogen Peroxide. Plant Cell Physiol. 42, 286–291. 10.1093/pcp/pce032

Picapietra, G., Acciaresi, H.A., 2022. OVERCOMING SEED DORMANCY OF JUNGLERICE (Echinochloa colona). Chil. J. Agric. Anim. Sci. 38, 154–163. 10.29393/CHJAA38-15OSGH20015

Priatama, R.A., Pervitasari, A.N., Park, S., Park, S.J., Lee, Y.K., 2022. Current Advancements in the Molecular Mechanism of Plasma Treatment for Seed Germination and Plant Growth. Int. J. Mol. Sci. 23, 4609. 10.3390/ijms23094609

Puglia, G.D., 2024. Reactive oxygen and nitrogen species (RONS) signalling in seed dormancy release, perception of environmental cues, and heat stress response. Plant Growth Regul. 103, 9–32. 10.1007/s10725-023-01094-x

Sajeev, N., Koornneef, M., Bentsink, L., 2024. A commitment for *life:* Decades of unraveling the molecular mechanisms behind seed dormancy and germination. Plant Cell 36, 1358–1376. 10.1093/plcell/koad328

Sivachandiran, L., Khacef, A., 2017. Enhanced seed germination and plant growth by atmospheric pressure cold air plasma: combined effect of seed and water treatment. RSC Adv. 7, 1822–1832. 10.1039/C6RA24762H

Tian, Z., Shen, G., Yuan, G., Song, K., Lu, J., Da, L., 2020. Effects of Echinochloa crusgalli and Cyperus difformis on yield and eco-economic thresholds of rice. J. Clean. Prod. 259, 120807. 10.1016/j.jclepro.2020.120807

Travlos, I., Gazoulis, I., Kanatas, P., Tsekoura, A., Zannopoulos, S., Papastylianou, P., 2020. Key Factors Affecting Weed Seeds’ Germination, Weed Emergence, and Their Possible Role for the Efficacy of False Seedbed Technique as Weed Management Practice. Front. Agron. 2, 1. 10.3389/fagro.2020.00001

Waskow, A., Howling, A., Furno, I., 2021. Mechanisms of Plasma-Seed Treatments as a Potential Seed Processing Technology. Front. Phys. 9, 617345. 10.3389/fphy.2021.617345

Wojtyla, Ł., Lechowska, K., Kubala, S., Garnczarska, M., 2016. Different Modes of Hydrogen Peroxide Action During Seed Germination. Front. Plant Sci. 7. 10.3389/fpls.2016.00066

Zhang, Z., Cao, J., Gu, T., Yang, X., Peng, Q., Bai, L., Li, Y., 2021. Co-planted barnyardgrass reduces rice yield by inhibiting plant above- and belowground-growth during post-heading stages. Crop J. 9, 1198–1207. 10.1016/j.cj.2020.10.011

Zwanenburg, B., Mwakaboko, A.S., Kannan, C., 2016. Suicidal germination for parasitic weed control. Pest Manag. Sci. 72, 2016–2025. 10.1002/ps.4222

